# Top-down feature-based cueing modulates conflict-specific ERP components in a Stroop-like task with equiprobable conditions

**DOI:** 10.1101/807412

**Authors:** Daniela Galashan, Julia Siemann, Manfred Herrmann

## Abstract

Both attention and interference processing involve the selection of relevant information from incoming signals. Studies already show that interference decreases when the target location is precued correctly using spatial cueing. Complementary, we examine the effect of feature-based attentional cueing on interference processing. We used a design with equal stimulus probabilities where no response preparation was possible in the valid condition due to a response mapping alternation from trial to trial. The color of the Stroop stimuli was precued either validly or invalidly. Electrophysiological data (EEG) from 20 human participants are reported. We expected reduced interference effects with valid cueing for behavioral data and for both Stroop-associated event-related potential (ERP) components (N450 and sustained positive potential; SP). The N450 showed a significant effect for valid trials but no effect in the invalid condition. In contrast, the SP was absent with valid cueing and present with invalid cues. These findings suggest that focused feature-based attention leads to a more effective attentional selectivity. Furthermore, the top-down influence of feature-based attention differentially affects the N450 and SP components.

## 1. Introduction

In this study the influence of feature-based attention on interference processing is investigated in a variant of the Stroop task with equiprobable stimuli. For this purpose, EEG data were recorded from human participants and the ERP components N450 and conflict SP were examined during processing of our Stroop-like task with feature-based attentional cueing (valid, invalid conditions).

In general, selective attention enables us to concentrate our limited processing resources on action relevant information while irrelevant information is suppressed. Thus, a cue stimulus improves performance when it correctly guides attention to an upcoming stimulus property (valid cueing; e.g. to a specific location or color). In return, performance deteriorates when attention is oriented to an incorrect stimulus property (invalid cueing; Posner, 1980). Selective attention leads to a more robust neuronal representation of attended and relevant incoming information compared to non-attended information (see Noudoost et al. 2010). Attentional orienting can happen either on purpose (top-down) or triggered by stimuli (bottom-up; Corbetta & Shulman 2002). Furthermore, the attentional focus can be directed to different stimulus properties: To a spatial position (spatial attention; Posner & Presti 1987), an object (Duncan 1984), or to non-spatial properties like color or movement (feature-based attention; Allport 1971). In the present task we used top-down feature-based cueing (valid, invalid) to orient attention to the feature ‘color’.

Interference processing influences performance and is closely related to selective attention. It is required when distracting task-irrelevant information impairs task performance. The task-irrelevant information influences task processing due to an overlap of this conflicting information with task-relevant stimulus or response dimensions (Kornblum, 1994). There are different types of interference effects (e.g. the flanker effect, Eriksen & Eriksen 1974; the Simon effect, Simon & Rudell 1967; the Stroop effect, Stroop 1935). For example, in the classical color-word Stroop task (Stroop 1935) the ink color of a color word has to be named. Thus, interference occurs when ink color and color word are different (incongruent condition) and there is no interference when both are identical (congruent condition). In the neutral condition non-color words are presented or letter strings like XXXX. The Stroop task has become a well-established paradigm in the study of interference (MacLeod 1991) and feature-based attentional mechanisms (Polk et al. 2008).

We used a Stroop task and recorded EEG (electroencephalogram) data to investigate subtle modulations in interference processing that might not show up in performance data. Two ERP (event-related potential) deflections are usually associated with Stroop interference processing: A N450 component and a sustained potential (SP; Larson et al. 2014) also called Incongruency Negativity (N_INC_; see Appelbaum et al. 2014). The N450 component is a relative negativity around 450ms for incongruent compared to congruent stimuli. Sometimes this component is labeled as medial frontal negativity (MFN, see Larson et al., 2014). The local distribution of the N450 can vary from fronto-central (Larson et al. 2014; Markela-Lerenc et al. 2004) to central and centro-parietal electrodes (Appelbaum et al. 2009; Chen et al. 2011). The function of the N450 is still under debate (Larson et al. 2014). Currently, it is most strongly associated with conflict detection processes (Hanslmayr et al. 2008; West et al. 2004; Larson et al. 2014). The N450 component appears to be involved in automatic monitoring processes but seems to be unaffected by top-down adaptation mechanisms (Larson et al. 2009). Other authors argue that the N450 constitutes a conglomerate of several subcomponents including one that is responsible for the top-down proactive adaptation between trials (Chen et al. 2011). The N450 amplitude is increased with fewer incongruent trials (e.g. West & Alain 2000; Tillman & Wiens 2011). The neural generator of the N450 is assumed to be located in ACC (e.g. Beldzik et al. 2015; Badzakova-Trajkov et al. 2009; Hanslmayr et al. 2008; Liotti et al. 2000).

The conflict SP (indicating “conflict sustained potential”, see West 2003; or “conflict slow potential”, see Larson et al. 2009) is a sustained relative positive potential for incongruent stimuli starting at about 500ms. In some publications the SP is denoted as “negative slow wave” (NSW, see Larson et al. 2014) or “late positive component” (LPC, see Liotti et al 2000). Its scalp distribution varies from temporo-parietal (West & Alain 1999; West & Alain 2000) to centro-parietal (Coderre et al. 2011), parietal (Markela-Lerenc et al. 2004; West 2003), and parieto-occipital sites (Appelbaum et al. 2009; Hanslmayr et al. 2008). The conflict SP is generally associated with conflict resolution and adjustment processes, though its exact functional role is still unclear (Larson et al. 2014). For example, it is involved in perceptual processing, but only for incongruent and neutral Stroop stimuli. Therefore, it might reflect the retrieval of the previously suppressed word information (West & Alain 2000). Contrary to the N450 the SP might also be an indicator of the cognitive control needed for a trial, as it is sensitive to previous trial congruency. Hence, it could also reflect neural mechanisms calling for increased attentional control (Larson et al. 2009). For the conflict SP, neural sources were reported in posterior brain regions as well as in bilateral frontal brain regions (Hanslmayr et al. 2008; West 2003).

When combining both influencing factors (selective attention and interference processing), it is interesting to examine how top-down attentional cueing affects interference processing. We focused on the influence of feature-based attention on Stroop interference expressed in modulations of the N450 and the SP components. Lupiáñez and Funes (2005) addressed this issue solely for behavioral data in a spatial Stroop task with spatial cueing. Interference effects in their task decreased when the stimulus location was validly cued, comparable to studies combining the flanker task with spatial cues (e.g. Callejas et al. 2004; McCarley & Mounts 2008; Munneke et al. 2008). However, Lupiáñez and Funes (2005) found interactions between interference processing and spatial cueing only for tasks with stimulus-stimulus conflicts and not with stimulus-response conflicts. As the Stroop task contains a stimulus-stimulus conflict (see Kornblum & Lee, 1995) it is a good candidate to investigate if this kind of interaction can also be found with feature-based cueing instead of spatial cueing and if this influence of attentional cueing is also expressed in neuronal processing. Concerning another stimulus-stimulus conflict (flanker effect) one study observed no interaction between feature-based cueing and flanker effects (Siemann et al. 2018). Nevertheless, it is interesting to examine Stroop effects with attentional cueing as Stroop conflict resolution appears to affect neuronal processing at a later stage during stimulus processing than flanker conflict resolution (Rey-Mermet et al. 2019). Furthermore, via dorsal-ventral brain connections focused attention (valid cues) might lead to inhibited processing of distracting information that is not related to the target (DiQuattro et al. 2014). To our knowledge there are no other studies on the influence of non-spatial attentional selection on Stroop interference processing. The Stroop task is especially suited for this research question as it might be susceptible to effects of attentional cueing due to its stimulus-stimulus conflict (see Lupiáñez & Funes 2005) and because relevant and irrelevant stimulus properties include the feature ‘color’ that can be used for feature-based cueing.

Summarized, in the present study we examined whether feature-based cueing of the upcoming stimulus color modulates task performance and neuronal processing reflected in behavioral data and Stroop conflict-specific ERP components (N450, conflict SP).

On a behavioral level valid cues were expected to induce an attenuation of the Stroop effect (difference: incompatible - compatible condition) compared to invalid cues. Assuming the N450 reflects conflict detection we expected an attenuated Stroop effect in the valid cueing condition as the correct cueing should lead to a facilitated stimulus processing. Concerning the conflict SP as an indicator of required cognitive control we hypothesize a higher Stroop effect with invalid cueing. In these cases additional control might be required to overcome biased expectancies.

## 2. Materials and Methods

### 2.1 Participants

EEG data were recorded from 23 volunteers (6 male; mean age = 23.6 years; SD = 2.9) without neurological or psychiatric history, substance abuse or medication affecting the central nervous system. All of them were native German speakers, had normal or corrected-to-normal vision, and were right-handed (assured by the Edinburgh inventory from Oldfield, 1971; median = 92.3 percent, range = 92.3 - 100 percent). Color blindness was tested with a modified version of the Ishihara Test for Colour Blindness (Ishihara, 1917) to ensure that all participants were able to discriminate colors. The experiment was approved by the local ethics committee and fulfilled the criteria of the Helsinki Declaration of the World Medical Association (Rickham 1964). Participants’ written informed consent was obtained from all individual participants included in the study prior to their participation, they were informed about the procedure of recording and their right to quit the experiment at any time. Moreover, they were assured that their data would be anonymized so that the possibility of backtracking of their identity from the data was excluded. They were paid 15 € or given course credit. Each participant underwent a training session and a separate EEG session. Three data sets had to be excluded from further analysis due to massive artifacts (excessive muscle artifacts; alpha waves dominating ERPs) leading to 20 analyzed and reported data sets.

### 2.2 Apparatus and Stimuli

The stimulus set was composed of four different colors and the respective color words. The experiment consisted of four congruent (CON) Stroop stimuli (each of the four color words in the congruent color), four neutral (NEU) Stroop stimuli (“xxxx” in any of the four colors) and 12 incongruent (INC) Stroop stimuli (each of the four color words in the remaining three incongruent colors). The frequency of all stimulus combinations was equal in all blocks and each congruency level appeared in 1/3 of all trials. Cue validity was 50%, i.e. in half of the cases the cue predicted the correct target color (valid) and in the other half an incorrect color (invalid). On invalid trials, the cue was never identical to the color word, thus avoiding additional interference due to cueing of the incorrect response. Usually, in most attentional cueing tasks higher stimulus frequencies are used for the valid cueing condition to ensure a benefit of attentional orienting. Instead, we used equal frequencies for all conditions because oddball stimuli with lower stimulus frequencies lead to expectation violation and to an engagement of specific brain networks (Kim 2014). Unfortunately, some of these brain regions influenced by differing stimulus probabilities (see e.g. Corbetta and Shulman, 2002; Vossel et al., 2006) are key regions also involved in attentional processing. Therefore, Macaluso and colleagues (2013) advise to use designs where it is possible to differentiate between effects of attention and effects of differing stimulus frequencies. An even stronger argument for the use of equal probabilities in the present study is the fact that several ERP components are influenced by varying stimulus probabilities (e.g. P2, P3, see Sutton et al. 1965; N450, see Lansbergen et al. 2007; Tillmann & Wiens 2011, West & Alain 2000). Thus, at least for the N450 in a design with unequal stimulus probabilities we would not be able to determine the actual attention and interference effects. Therefore, to disentangle effects of selective attention and Stroop interference from effects purely elicited by differing stimulus frequencies, we used equal probabilities for all cueing conditions (each ½ valid/ invalid) and for all Stroop conflict conditions (each 1/3 compatible/ incompatible/ neutral). To ensure participants’ compliance with our instructions and a correct attentional orienting despite the 50% cueing validity we used carefully designed task instructions (see below). Furthermore, we knew from other studies that a comparable approach with 50% cue validity is indeed working and participants are able to follow the instructions (Siemann et al. 2016; Galashan & Siemann 2017).

The combination of two validity levels (valid/invalid) and three congruency levels (CON/INC/NEU) resulted in six conditions appearing with equal frequencies in 1/6 of all trials.

We controlled for sequential effects (conflict adaptation effects, see Gratton et al. 1992) as these effects influence the conflict SP component (Larson et al. 2009, 2012). Therefore, each experimental block consisted of 144 pseudo-randomly distributed trials (24 trials/condition) with a fixed pseudo-randomized order ensuring that each of the six conditions was followed equally often by each of the others within each block. Block order was counter-balanced over participants. All participants completed three training blocks in the training session and six experimental blocks with short breaks of a few minutes in between in the EEG session. The parameters of the training session matched those of the experiment apart from the first training block where written feedback was provided following every trial (“correct“, “incorrect“, or “too slow” for responses > 1000ms after stimulus onset).

Stimulus presentation was based on the Presentation®-Software (Neurobehavioral Systems; https://nbs.neuro-bs.com) using a PC connected to a 19 inch computer monitor (viewing distance = 56cm). A smoothed fixation point (1.23° × 1.23°) was presented at the beginning of a trial and after cue presentation. The cue word appeared in white letters and could be either of the German words “rot” (red, 1.74° × 1.02°), “blau”(blue, 2.66° × 1.23°), “gelb” (yellow, 2.61° × 1.54°) or “grün” (green, 2.66° × 1.28), as well as “xxxx” (3.17° × 0.72°) for the NEU condition. Cues and stimuli appeared centrally in lower case (Times New Roman). The common fixed response mapping (e.g. red, green, blue, yellow each assigned to one of the 4 response fingers) would allow for a complete response preparation before stimulus preparation in all validly cued trials. In contrast, all invalidly cued trials would require a switch from the prepared response to another response finger. Thus, correct/ incorrect response preparation would have been confounded with cueing. In order to avoid this confound and prevent motor preparation based on cue information, the colors were reassigned to the respective response buttons on the computer keyboard on every trial (see e.g. Fehr et al., 2007). Therefore, the keys-to-color allocation was variable and changed from trial to trial. For that purpose four equal-sized boxes (1.4° × 2.9°) arranged horizontally were presented below the stimulus (y=−1.4°) with a small gap in between (0.1°, see Figure 1). The array of boxes extended 5.9° to each side. The boxes were filled with the four colors and corresponded to the four response keys on the computer keyboard. Responses were given with four fingers (left and right index and middle fingers) on the computer keyboard (adjacent keys f, g, h, and j).

**Figure 1:**
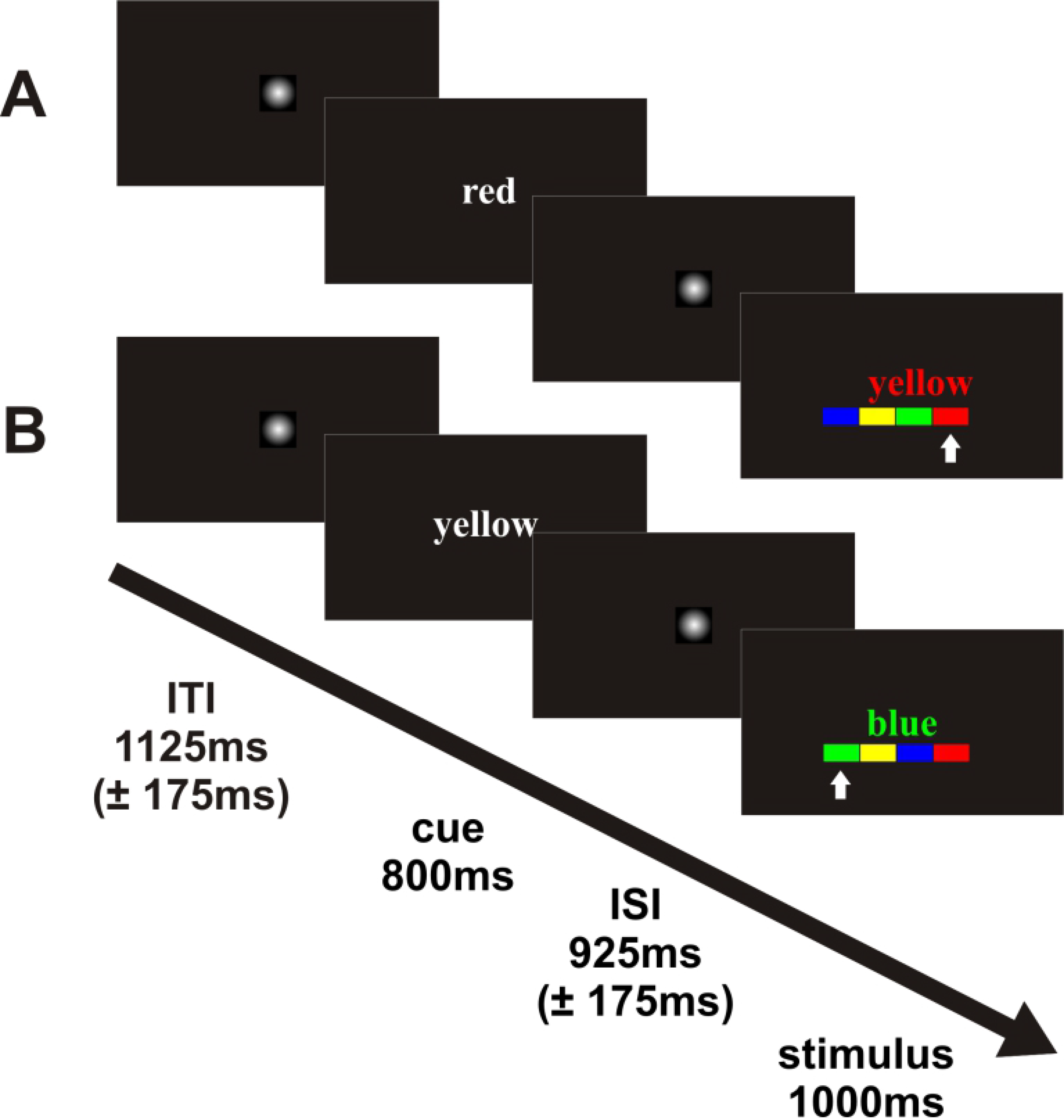
Sample trials with time course. After the ISI the color word appeared together with the colored boxes and cued the to-be-attended color. Participants had to press the button on the keyboard corresponding to the box below the stimulus with the appropriate color (indicated by white arrows in the figure). A) valid cueing; B) invalid cueing; all stimuli presented here are incongruent Stroop stimuli.

Participants were instructed to use the cue information and to direct their attention to the corresponding color and to hold fixation at the center of the screen without overt eye movements to the colored boxes below. In the instruction, particular emphasis was placed on the fact that they had to respond as fast and correct as possible especially on trials with correct cueing. They were informed that they needed to orient their attention according to the cue to achieve this goal. Other studies also confirm that this approach works even with 50% cue validity (Siemann et al. 2016; Galashan & Siemann 2017). Furthermore, they were asked to avoid eye and head movements. Each trial started with a jittered intertrial interval (ITI) showing a fixation point (1125ms ± 175ms). Thereafter, the cue word appeared for 800ms, followed by a jittered interstimulus interval (ISI) where the fixation point reappeared (925 ±175ms). Finally, the stimulus and the colored boxes were presented simultaneously for 1000ms (see Figure 1).

All participants completed two separate sessions. First, they were familiarized with the experimental design and the response procedure in a training session using a standardized written task instruction. Participants were informed about the probability of the cue validities and it was emphasized that it is important to follow the cue and correctly orient attention to the upcoming color in order to benefit from the cueing information in cases of a valid cue. The data of the training session were analyzed to check if participants showed an attention effect (invalid - valid cueing) to make sure they understood and followed the instruction despite equal cue validities. They were instructed to perform at high speed while simultaneously taking care to keep the error rate low. The experimental session with EEG measurements took place on a separate day (average time period between training and EEG session: 3.5 days; SD = 3.2). Before EEG recording the first five trials of a no-feedback training block were presented to remind the participants of the task requirements. Training and experiment were conducted in a dimly lit room where participants sat on a height adjustable swivel chair in front of a computer screen. They were asked to position their head on a chin rest to ensure a fixed distance from monitor and to minimize head movement.

### 2.3 Data Acquisition

EEG was recorded from 64 channels (Neurofax μ electroencephalograph (Nihon Kohden; Tokyo, Japan) with the program eemagine EEG 3.3 (Medical Imaging Solutions GmbH). The REFA® multichannel system (TMS International; www.tmsi.com) served as a direct-coupled (DC) amplifier (sampling rate: 512 Hz). Data were average-referenced and impedances were kept below 5kΩ. An array of 64 Ag/AgCl sintered ring electrodes was arranged according to the extended international 10-20 system using a standard recording cap (EASYCAP, www.easycap.de). The ground electrode was placed at the left mouth angle and four additional electrodes (infra- and supraorbitally to the right eye and at the outer canthi) were used for recording of the electrooculogram (EOG).

### 2.4 Data Analysis

#### 2.4.1 Behavioral data

Behavioral data were analyzed with IBM SPSS Statistics for Windows (Version 11.5, SPSS Incorporation). After confirmation of normal distribution (Kolmogorov-Smirnov-Test), mean reaction times (RTs) averaged over all six blocks and each participant per condition were fed into a validity (2: valid and invalid) × congruency (3: congruent, incongruent and neutral) repeated measures analysis of variance (rmANOVA). A standard significance level of α=0.05 was adopted. If necessary, Greenhouse Geisser corrected epsilon values were used whenever sphericity could not be presumed (as assessed by Mauchly’s Test). Only correct trials were included in the analysis of RTs. Significant effects were investigated post-hoc using paired t-tests. For visualization of the RTs, different effects were computed: The Stroop effect (incompatible - compatible condition), the facilitation effect (compatible vs. neutral), and the interference effect (incompatible vs. neutral). Error rates (percentage of incorrect trials and misses per condition) were analyzed separately. As not all error rate data were normally distributed a non-parametric Friedman Test was performed and post-hoc comparisons were carried out with Wilcoxon Tests.

#### 2.4.2 EEG data

EEG data were analyzed with BESA® 2000 (Brain Electrical Source Analysis; MEGIS Software GmbH, 2002, www.besa.de) using a low-cutoff filter (0.03 Hz forward 6db/oct) and a notch filter (50Hz, width 2). After visual inspection single channels were interpolated or defined as ‘bad channels’ when necessary (0 – 6 altogether per participant; mean= 2.4). Only in one case an electrode channel used for the subsequent analyses (F4) had to be interpolated and no analyzed channel had been defined as a ‘bad’ channel. Afterwards, EEG epochs of correct trials were averaged stimulus-locked to the target stimulus from −400ms to 1000ms. Erroneous and artifact-contaminated trials (exceeding 100μV, with eye blinks, excessive eye movements or muscle activity) were excluded from further analyses. Due to differing error rates there were more trials for compatible conditions. To ensure comparable averages we aimed at identical trial numbers for all averaged conditions. Thus, of the remaining trials exactly 100 trials per condition and participant were averaged based on visual inspection. The trials were selected to be uniformly distributed over the whole length of the experiment to avoid a bias towards a specific phase of the experiment (start, end). We performed statistical analyses on mean amplitude values of each component using repeated measures ANOVAs with the factors *validity* (valid, invalid cueing) and *congruency* (congruent, incongruent, neutral Stroop stimulus)for both ERP components. Additionally we captured the topographical distribution with component-specific levels of the factors *frontality* and *laterality*: In case of the N450 (electrodes: F3, Fz, F4, C3, Cz, C4, P3, Pz, P4), there were three frontality levels (F, C, P) and three laterality levels (3, z, 4). For the SP (electrodes C1, Cz, C2, P1, Pz, and P2), we us two frontality levels (C, P) and three laterality levels (1, z, 2).The interaction of cue validity and Stroop congruency was further investigated using post-hoc paired t-tests with a Bonferroni-Holm-corrected threshold (based on α <.05). The N450 component was measured between 300 and 430ms, the time window for the SP was 700-800ms.

## 3. Results

### 3.1 Behavioral Data

Figure 2 shows the Stroop (incongruent - congruent), facilitation (compatible - neutral), and interference effects (incompatible - neutral) of mean reaction times (RTs) and error rates. The repeated measures ANOVA on the RTs with validity (valid and invalid) and congruency (congruent, incongruent, neutral) yielded only main effects for the validity and congruency effects respectively (*F*_(1;19)_ = 52.98, p < .001 and *F*_(1.4;27)_ = 49.02, p < .001), whereas the interaction effect of these factors was not significant (validity x Stroop: *F*_(2;38)_ = 1.5, p = .227).

**Figure 2:**
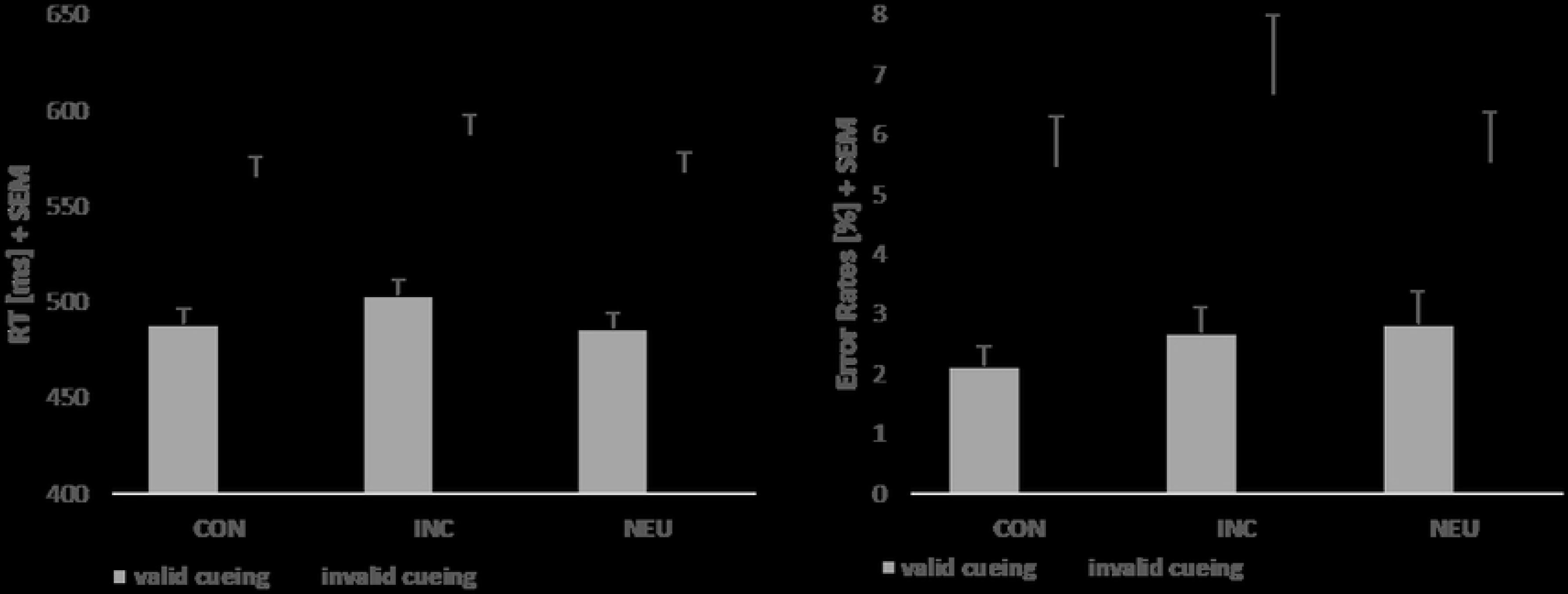
Reaction times (RTs) and error rates. RTs (left, in ms) and percentage error rates (right, in %) with valid (grey bars) and invalid cueing (black bars), with 1 Standard Error of the Mean (SEM); CON = congruent Stroop stimulus, INC = incongruent Stroop stimulus, NEU = neutral Stroop stimulus

The post-hoc analyses for RT data showed that participants were significantly slower at responding to incongruent than congruent (*t*_(19)_=6.8; p<001; mean Stroop effect: 18.3ms; SD: 11.9; Cohen’s d = 0.4) and neutral stimuli (interference effect; *t*_(19)_=8.4; p<.001; mean interference effect: 19ms; SD: 10.1; Cohen’s d = 0.4). Conditions did not significantly differ for the facilitation comparison (neutral vs. congruent, *t*_(19)_=−0.523, p=.607). Concerning the attentional cueing manipulation, RTs were faster for valid than invalid trials (*t*_(19)_=7.3; p<.001; mean attention effect: 83.2ms; SD: 51.1; Cohen’s d = 1.4).

Analyzing the error rates revealed a main effect of cue validity (*F*_(1;19)_ = 15.6, p < .005) with more errors evident on invalidly cued trials compared to valid cueing (*t*_(19)_= 3.9; p<.005; Cohen’s d = 0.7). The Stroop main effect (*F*_(1.5;29.3)_ = 1.577, p = .225) as well as the interaction (validity × Stroop: *F*_(2;38)_ = 3.065, p = .58) were not significant. Post-hoc comparisons also yielded non-significant results (incongruent vs. congruent: *t*_(19)_= 1.482; p = .155; neutral vs. congruent: *t*_(19)_= .861; p = .4; incongruent vs. neutral: *t*_(19)_= 1.081; p = .293).

### 3.2 EEG Data

As illustrated in Figure 3, visible differences between incompatible and compatible conditions occurred relatively early with valid cueing (left), presumably covering the N450 time window and with invalid cueing later in the SP time window (right).

**Figure 3:**
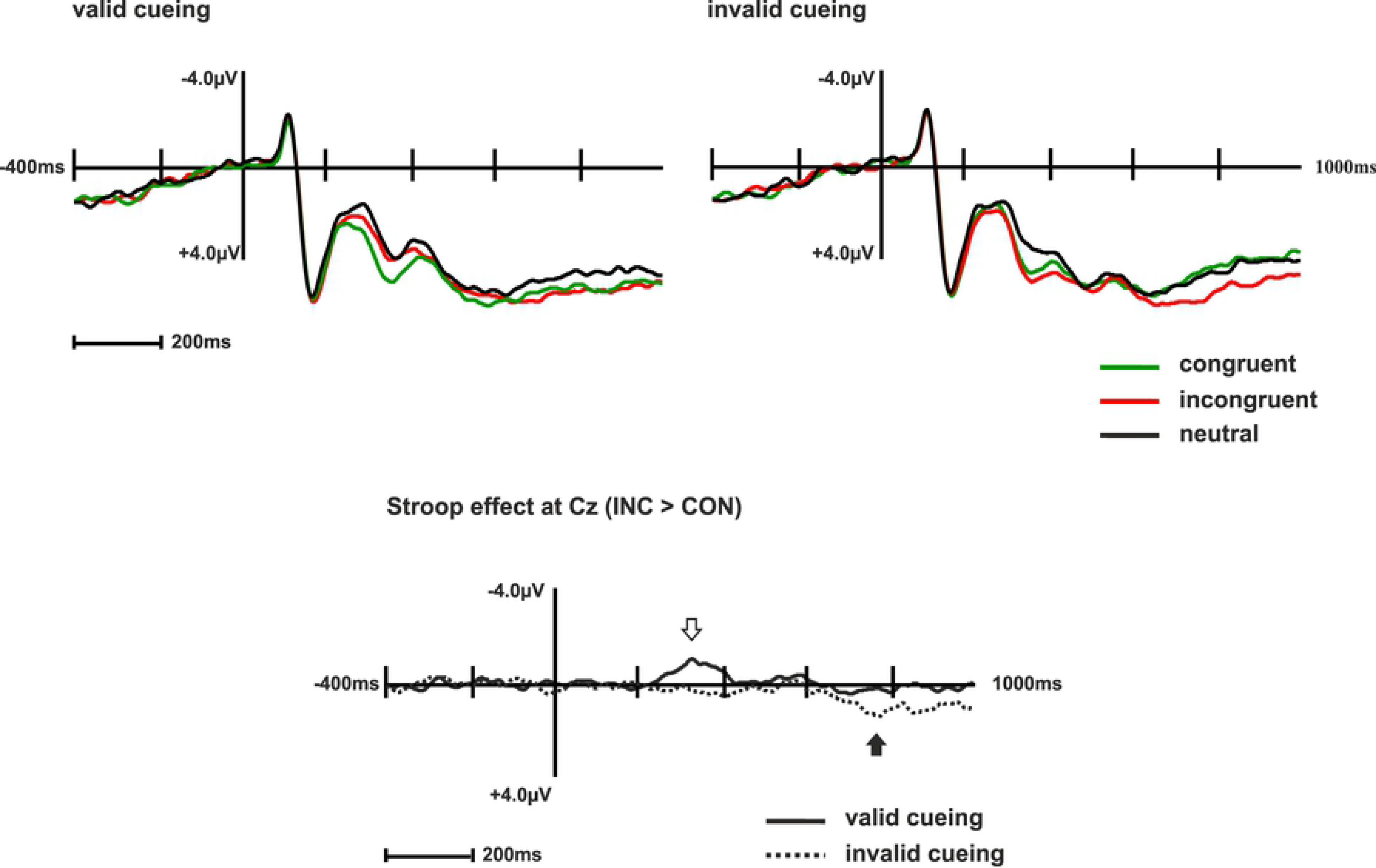
ERPs and difference waves. Top: ERPs (stimulus-locked grand average: -400ms to 1000ms) of congruent (green), incongruent (red) and neutral stimuli (black line) with valid (left) and invalid (right) cueing at electrode position Cz. Bottom: difference wave of the Stroop effect (incongruent-congruent) with valid (solid line) and invalid cues (dashed line) at electrode Cz. Arrows indicate the N450 and the SP components (blank = N450; filled = SP).

ERP data were analyzed using repeated-measures ANOVAS with the factors validity (valid, invalid) and congruency (congruent, incongruent, neutral). To capture the topographical distribution, the factors frontality and laterality were as follows for the respective ERP components: N450 (frontality levels F, C, P; laterality levels 3, z, 4); SP (frontality levels C, P; laterality levels 1, z, 2).

The analysis yielded a main effect of Stroop congruency (*F*_(2;38)_= 10.9; p<.0.001) as well as cue validity during the N450 time window (*F*_(1;19)_= 9.5; p<.01).These factors additionally interacted with each other (*F*_(2;38)_= 11.7; p<.001). Post-hoc paired t-tests showed that the Stroop effect was significant after valid (*t*_(19)_=5.7; p<.001) but not after invalid cueing (*t*_(19)_= −0.9; p>.3). The SP was also characterized by main effects of Stroop congruency (*F*_(2;38)_= 5; p<.05), cue validity (*F*_(2;38)_= 7.2; p<.05), as well as an interaction of both factors (*F*_(2;38)_= 5.1; p<.05). Post-hoc analyses yielded a significant Stroop effect after invalid cues only (*t*_(19)_= −0.5; p>.6 and *t*_(19)_= −3.6; p<.005 for valid and invalid cueing respectively).

Concerning the topographic distribution, the N450 showed an interaction effect of Stroop congruency and validity with the factor laterality (*F*_(4;76)_= 2.6; p<.05). No such effects were apparent for the SP time window.

Figure 4 illustrates the scalp distributions of the significant Stroop effects (incongruent – congruent) in the N450 and SP time windows (correspondingly for valid and invalid cueing).

**Figure 4:**
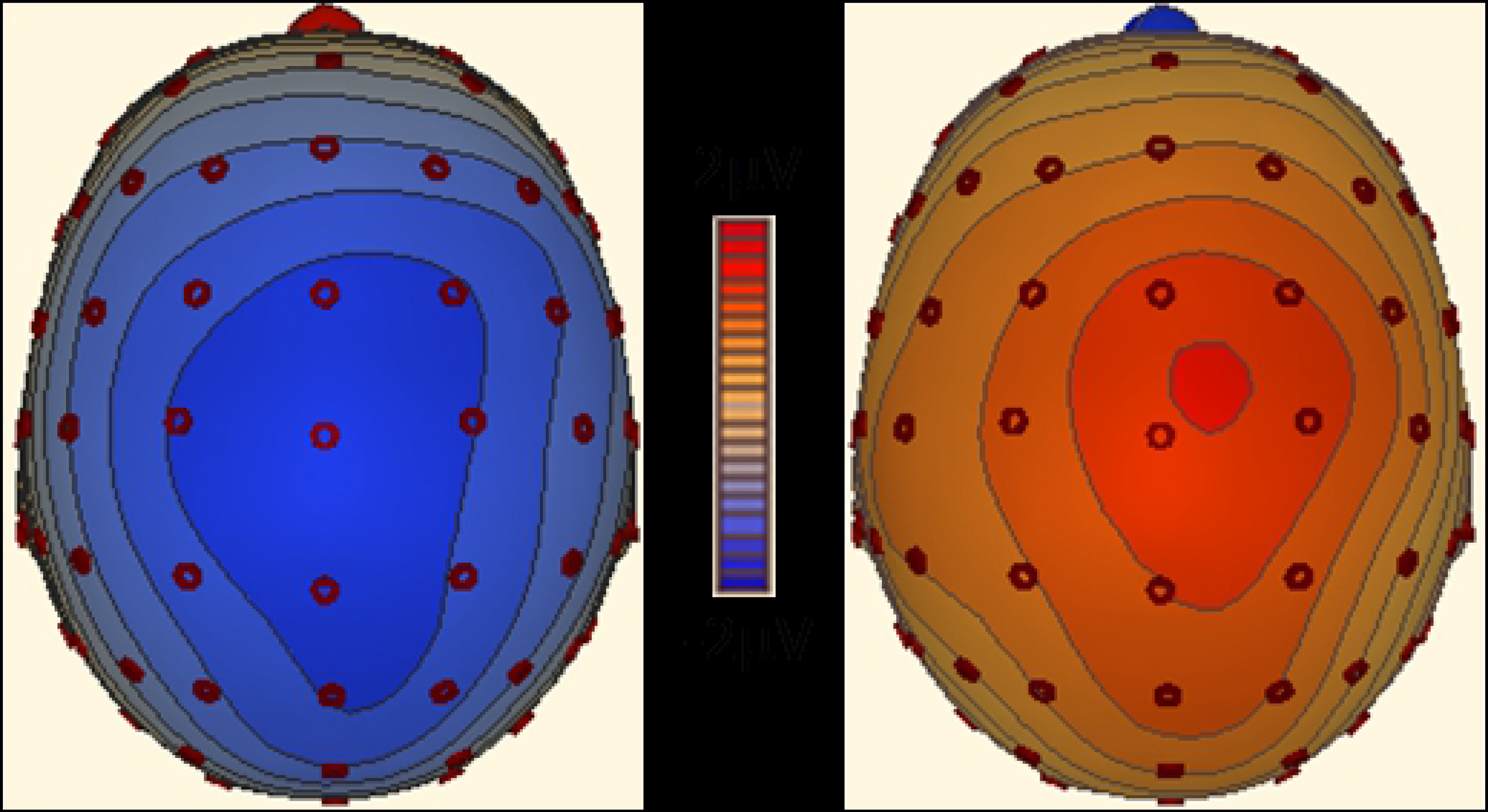
Stroop conflict shown with topographic maps. Topographic maps of the Stroop effect (incongruent - congruent) during the N450 time window (365ms) with valid cueing (left) and during the SP time window (773ms) with invalid cueing (right).

## 4. Discussion

In order to examine the influence of feature-based attention on interference processing, we combined attentional cueing with Stroop conflict processing and assessed cueing effects in the ERPs during the N450 and SP time windows.

The behavioral data analysis shows that both effects of interest (Stroop effect and attention effect) were elicited by the present experimental design. However, the cueing manipulation did not significantly reduce the amount of Stroop interference when cues were valid. Hence, our first assumption (attenuation of the Stroop effect with valid cueing) could not be confirmed. Nevertheless, the observed main effects pointed to the fact that a Stroop effect was present, and the cueing manipulation worked despite equal probabilities for valid and invalid cues. The lacking facilitation effect (congruent vs. neutral) in the RTs (see Figure 2) indicates that the differences in processing between our neutral and congruent Stroop conditions did not turn out as expected. RTs for neutral and congruent conditions were almost identical (valid cueing: 481 vs. 484ms; invalid cueing: 565 vs. 563ms). With this result a cautious interpretation of the neutral condition in the present design is warranted as its RT values do not lie in between congruent and incongruent conditions and it even shows the highest error rates with valid cueing. However, other Stroop studies also yield comparable RTs for neutral and congruent conditions (e.g. West & Alain 2000) and the processing of the neutral conditions is assumed to depend on the choice of the neutral stimuli (shapes, non-word syllables, color-neutral words…) and their probability (see e.g. Shichel & Tzelgov 2017).

The ERP data shed light on differences between the cueing conditions that were not visible in the behavioral data. The Stroop effect shows validity-specific modulations for both ERP components, as we found substantial differences between the effects of valid and invalid cueing on Stroop stimulus processing in both analyzed time windows. On the one hand, a relative negativity during the incongruent condition in the N450 time window was only observable with valid cueing. This finding contradicts our hypothesis of facilitated stimulus processing leading to reduced conflict detection in the N450 time interval. On the other hand, a relative positivity of incongruent compared with congruent stimuli (Stroop effect) was specific for invalid cueing within the SP interval. This result confirms our assumption of an increased demand for cognitive control reflected in a higher Stroop effect with invalid cueing in the SP time window and a corresponding reduction with valid cueing.

In the following, we initially discuss the effects visible in the N450 time window and then the effects in the conflict SP interval followed by possible explanations holding for both effects. All discussed effects refer to the Stroop comparison (congruent vs. incongruent), and references to the associated facilitation (congruent vs. neutral) and interference (incongruent vs. neutral) components of the Stroop effect are explicitly stated where applicable.

The fact that there was a significant Stroop effect in the N450 interval with valid cueing even though it was not present on invalidly cued trials contradicts the hypothesized reduction of conflict detection with valid cueing. Still, in a digit version of the Stroop task with a comparable cueing manipulation the N450 was also shown to be present in the valid cueing condition (Szűcs & Soltész 2012). In their experiment the N450 was visible in the valid condition although they demonstrated that valid cueing effectively reduced response conflict (using electro-myography/ EMG and the lateralized readiness potential/ LRP). This led to the suggestion that the N450 might not be involved in response conflict processing (see also Mager et al. 2007). Indeed, with their valid cueing a complete response preparation was possible as they used a fixed stimulus-response mapping and effective cues had an accuracy of 100 percent. This might have led to confounding response preparation and cueing in their study. Thus, our study adds the finding that a N450 can also be present with valid cueing in cases when no response preparation is possible.

However, on invalidly cued trials no N450 differences between congruent and incongruent conditions were detectable. This finding might be attributed to the higher attentional demands evoked by invalidly cued trials. When cues were incorrect, attention had to be redirected towards the actually presented color. This process might have added another source of conflict resulting in a disruption of the usual conflict detection processing in the Stroop task. Thus, it might be possible that the N450 component was masked by the identification of the invalid cue information. Incongruent Stroop stimuli require highly demanding information processing involving the suppression of conceptual level processing when the word deviates from the color (West et al. 2005). We suppose that only after having reset the goal enough resources were available to process the semantic meaning of the color word and thereby detect the Stroop conflict.

Another explanation for the lacking hypothesized decrease of interference with valid cueing might be the argument that the N450 is robust against top-down modulations. For example, Larson et al. (2009) demonstrated that the N450 is not correlated with behavioral indices of top-down conflict adaptation evoked by repetition effects. The authors concluded that the N450 is associated with automatic conflict monitoring. Likewise, in a study of Xiang et al. (2013) the N450 was evoked even though the color word was masked and therefore not consciously processed. The authors argue that the N450 represents a preconscious bottom-up process whereas the SP component is subject to voluntary manipulations and thus prone to top-down influences. Similarly, in stimulus onset asynchrony (SOA) studies several authors found an N450 in positive SOA conditions where the color appears shortly before the word (Appelbaum et al. 2012; Coderre et al. 2011). At the same time, the fact that the Stroop effect during the N450 time window in our study differed between cueing conditions contradicts these assumptions of automatic bottom-up processing insusceptible to top-down effects. Our data rather point to top-down influences on neuronal processing during this relatively early interval even if the hypothesized effects were observed in the other condition as expected. Notably, in the present study Stroop-related ERP differences under valid cueing were most pronounced between 250ms-350ms and terminated before 450ms (stimulus-locked). Such a temporal shift of the N450 is also observable with positive SOAs (Appelbaum et al. 2009; Coderre et al. 2011), leading to the assumption that the response is already in progress at the occurrence of the N450. A frequent finding with positive SOAs is also the lack of an SP component despite the presence of an N450 indicating that the conflict was detected (Coderre et al. 2011). Correspondingly, in the present study in contrast to a reliably evoked N450 no Stroop-related SP component was visible on validly cued trials. Coderre and colleagues (2011) propose that the SP is only present in those conditions where conflict resolution is required. This interpretation supports the initial hypothesis of the present study that valid cueing attenuates the Stroop effect. According to this line of argumentation, no SP component was observable because no conflict resolution was required with valid cueing. The conflict was detected (N450 modulation) but did not induce response activation. Similarly, Lamers et al. (2010) found that cueing two possible colors of an upcoming Stroop stimulus leads to increased attentional focusing on the eligible responses for that trial rather than inhibition of the uncued and therefore ineligible responses. These data further substantiate the assumption that top-down control is accompanied by enhanced selection mechanisms rather than active inhibition.

In contrast to the lacking Stroop effects with valid cueing in the SP time window we observed strong Stroop effects with invalid cueing. These findings are corroborated by the scalp distribution of the interference effect (incongruent vs. neutral) in the SP time window. It shows a relative negativity at posterior left-sided electrode sites of incongruent compared to neutral stimuli on invalidly cued trials. This topography is in line with a suggested re-activation of left-sided regions for incongruent words reflected in the SP (Liotti et al. 2000). Therefore, our findings substantiate the hypothesis that only invalidly cued incongruent stimuli caused suppression mechanisms of the word meaning and that this suppression was followed by reactivation processes. By contrast, valid cueing was characterized by conflict detection without suppression.

Two factors might have influenced the results for both ERP components. First, the variable response button allocation and second the frequency balanced task design. The alternating response mapping probably created an additional task demand that might have increased the overall task difficulty and could have occupied further information processing resources. Task difficulty is known to influence ACC activity (Paus et al. 1998), the brain structure assumed to be involved in the generation of the N450 component (e.g. Liotti et al. 2000). The flexibly changing response mapping possibly also led to reduced stimulus-response conflict compared to a fixed allocation where the word can more easily activate the response. Concerning the N450, there is no consensus about its presence solely based on a stimulus-stimulus conflict. While West et al. (2004) showed that a pure stimulus-stimulus conflict can suffice to elicit an N450, the data of Chen and colleagues (2011) contradict this view. A study by Donohue et al. (2016) indicates that the N450 is influenced when more response options are used. In the present design the changing mapping complicated the choice between response options even though there was a 1:1 mapping between the number of buttons and corresponding colors.

One might wonder if the alternating response boxes might have modified task processing in a way that participants only relied on cueing information and performed a simple match between the cued color and the response box instead of processing the target stimulus. Of course, for valid conditions this strategy would have led to fast RTs and low error rates. Additionally, there would be no Stroop effect with valid cueing as the target stimulus with its conflicting information would not have been processed. In turn, for invalid trials error rates and RTs would have considerably increased. The rather low error rates (5.7-7%) and not unusually long RTs (mean 563-584ms) in the invalid conditions as well as a comparable Stroop effect for valid and invalid conditions are arguments against this concern (difference of 5.4ms between valid and invalid Stroop effects). Furthermore, to prevent direct priming effects from the cue to the target display we used top-down cues without colors (white color words). Considering the data we assume the participants did not use a simple matching strategy and indeed processed the target stimulus as indicated by the existing Stroop effect also for valid trials. Nevertheless, we have to admit that a possible facilitation of the response button selection with attentional cueing instead of a facilitation of the target stimulus can’t be ruled out with the present design.

The second influencing factor, a frequency-balanced design, was needed to rule out potential probability effects. Nevertheless, it might have produced other modifications in task processing. The main effect of validity in the behavioral data disproves the possible concern that the attentional cueing was not followed due to the low predictive power of the cues. Participants indeed used cue information and oriented attention according to the cue.

Concerning stimulus frequencies in interference tasks, in designs with less frequent incongruent stimuli the level of conflict on incongruent trials is higher and the oddball effect additionally influences neuronal processing (see Kim, 2014 for a meta-analysis on the oddball effect). Tillman and Wiens (2011) only found a significant Stroop effect in the N450 time window when incongruent stimuli were rare (20%) but no effect with frequent incongruent stimuli (80%). Similarly, Lansbergen and colleagues (2007) reported higher amplitudes for the N450 and the conflict SP in their high conflict condition (20% incongruent) compared to the low conflict condition (80% incongruent). Another study (Chouiter et al. 2014) detected different brain sources associated with high and low levels of Stroop conflict although they defined high/low conflict in another way. As we also used neutral stimuli only 1/3 of our trials were incongruent but the frequencies of congruent and incongruent trials were identical in the present design. Nevertheless, it is likely that our frequency choice led to an attenuated Stroop effect in general for both analyzed ERP components.

West and Alain (2000) demonstrated that conceptual level information is generally suppressed by default when incongruent stimuli are frequent. Only when incongruent stimuli are rare conceptual information is the primary source of information. This frequency-specific effect primarily affects congruent stimuli because conceptual information of the word can guide responses only if color and word match. Therefore, when perceptual level processing is encouraged instead of conceptual level processing the N450 is possibly less pronounced in general. Furthermore, it is likely that on invalidly cued incongruent trials (where the perceptual information predicted by the cue was incorrect) participants reduced the attentional bias towards either perceptual or conceptual level information. This altered processing mode probably further reduced the N450 on invalidly cued trials when compared to classical Stroop studies where the congruent condition is usually presented more frequently.

It would be helpful to conduct further studies to disentangle effects caused by differing stimulus frequencies and by different types of response mappings on Stroop interference. Furthermore, it has to be investigated under which circumstances the N450 is prone to top-down influences as in the present study because former studies rather showed its insensitivity to top-down influences.

## 5. Conclusion

The present data show a significant Stroop effect in the N450 time window with valid but not with invalid cueing. Thus, contrary to other studies the N450 component was affected by top-down influences. We assume that attentional selection started relatively early during valid cueing conditions. Here, conflict detection might have taken place during the N450 time window.

By contrast, the SP component was absent with valid cueing. Thus, no conflict resolution mechanisms seemed to be evoked during this SP interval because the conflict detection (accompanied by the N450 component) did not lead to interference resolution. With invalid cueing, additional task-related processing mechanisms probably masked conflict detection and might explain the absent N450. Moreover, the incongruent word activated an incorrect response evoking conflict resolution and control mechanisms and leading to an SP. In sum, according to our results the N450 is involved in automatic conflict detection and may be influenced by enhanced attentional selection but not by active inhibition. By contrast, the SP reflects Stroop conflict resolution processes that are reduced or even absent with correct attentional direction to the upcoming stimulus color. Overall, the present study supports the idea that feature-based attention modulates interference processing. Furthermore, the top-down influence of attentional cueing differs for the N450 and the SP component.

## Acknowledgments

We would like to thank Dr. Thorsten Fehr for helpful advice concerning the alternating response mapping and EEG data analysis. Furthermore, we are grateful to Dr. Detlef Wegener for suggestions concerning the underlying idea of the study. Finally, we would like to thank our participants.

